# Multi-locus transcranial magnetic stimulation system for electronically targeted brain stimulation

**DOI:** 10.1101/2021.09.20.461045

**Authors:** Jaakko O. Nieminen, Heikki Sinisalo, Victor H. Souza, Mikko Malmi, Mikhail Yuryev, Aino E. Tervo, Matti Stenroos, Diego Milardovich, Juuso T. Korhonen, Lari M. Koponen, Risto J. Ilmoniemi

**Affiliations:** Department of Neuroscience and Biomedical Engineering, Aalto University School of Science, Espoo, Finland; BioMag Laboratory, HUS Medical Imaging Center, University of Helsinki and Helsinki University Hospital, Helsinki, Finland; School of Physiotherapy, Federal University of Juiz de Fora, Juiz de Fora, MG, Brazil; AMI Centre, Aalto NeuroImaging, Aalto University School of Science, Espoo, Finland; Institute for Microelectronics, Technische Universität Wien, Vienna, Austria

**Author notes:** Corresponding author; Postal address: Department of Neuroscience and Biomedical Engineering, Aalto University School of Science, P.O. Box 12200, FI-00076 AALTO, Finland. Contributed equally (Jaakko O. Nieminen, Heikki Sinisalo).

**Keywords:** Transcranial magnetic stimulation, mTMS, Multi-locus, Transducer, Coil, Electric field, Motor mapping

## Abstract

**Background:** Transcranial magnetic stimulation (TMS) allows non-invasive stimulation of the cortex. In multi-locus TMS (mTMS), the stimulating electric field (E-field) is controlled electronically without coil movement by adjusting currents in the coils of a transducer.

**Objective:** To develop an mTMS system that allows adjusting the location and orientation of the E-field maximum within a cortical region.

**Methods:** We designed and manufactured a planar 5-coil mTMS transducer to allow controlling the maximum of the induced E-field within a cortical region approximately 30 mm in diameter. We developed electronics with a design consisting of independently controlled H-bridge circuits to drive up to six TMS coils. To control the hardware, we programmed software that runs on a field-programmable gate array and a computer. To induce the desired E-field in the cortex, we developed an optimization method to calculate the currents needed in the coils. We characterized the mTMS system and conducted a proof-of-concept motor-mapping experiment on a healthy volunteer. In the motor mapping, we kept the transducer placement fixed while electronically shifting the E-field maximum on the precentral gyrus and measuring electromyography from the contralateral hand.

**Results:** The transducer consists of an oval coil, two figure-of-eight coils, and two four-leaf-clover coils stacked on top of each other. The technical characterization indicated that the mTMS system performs as designed. The measured motor evoked potential amplitudes varied consistently as a function of the location of the E-field maximum.

**Conclusion:** The developed mTMS system enables electronically targeted brain stimulation within a cortical region.

## Introduction

Transcranial magnetic stimulation (TMS) offers means to stimulate a specific cortical region non-invasively [1]. Since its first demonstration in the 1980s with a round coil [2], figure-of-eight coils [3] have become common, as they allow targeting TMS in a more specific manner. To adjust the stimulated cortical location, a TMS coil is typically moved manually. Robotic TMS systems offer an alternative approach [4–6]; however, the mechanical coil movement is relatively slow due to inertia and safety limitations. Thus, with a single-coil TMS system, it is practically impossible to adjust the stimulated spot fast, in the neuronally meaningful, millisecond timescale. With a pair of separate coils, it is also difficult to stimulate distinct nearby targets due to the relatively large coil size [7]. Fast stimulation of multiple cortical sites would enable the study of causal interactions in functional networks and more accurate and personalized treatments for neurological disorders [8]. There is a need for a flexible approach that allows TMS targeting based on real-time physiological feedback and convenient stimulation of nearby targets.

To overcome the slow mechanical coil adjustment and to enable the fast stimulation of different nodes of functional networks, we introduced multi-locus TMS (mTMS), in which a single coil is substituted with a transducer consisting of several coils [9]. By adjusting the relative currents in the coils, the induced electric field (E-field) pattern in the cortex can be modified electronically without coil movement. With mTMS, distinct cortical targets can be stimulated with sub-millisecond interstimulus intervals (ISIs) [10] and physiological feedback can be utilized in a closed loop to automate stimulation protocols [11,12]. Others have also taken steps towards implementing multi-coil TMS [13], with Navarro de Lara et al. presenting a prototype concept based on a 3-axis coil [14].

In this work, we aimed to develop an mTMS system that allows the adjustment of the location and orientation of the E-field maximum within a 2-dimensional (2D) cortical region 30 mm in diameter. Such a system would provide a substantial improvement on the 1-dimensional linear control that we previously achieved with a 2-coil mTMS system [9,15]. Here, we present our new mTMS system and demonstrate its unique capabilities in the context of automatic mapping of the primary motor cortex. The mTMS transducer developed in this study is based on the 5-coil concept we presented in [9].

## Methods

In this section, we introduce the key components of the mTMS system, i.e., electronics and the transducer. We also present the measurement protocols and analysis methods used to validate the system.

### Electronics

The mTMS system is based on independently controlled H-bridge circuits (Fig. 1B) [9,16–19]. The electronics can be roughly categorized into the following modules (Fig. 1A): control unit, charging unit, channels, coils, and auxiliary electronics. The control unit is responsible for the low-level control and operation of the system, whereas the charging unit, the channels, and the coils constitute the stimulation-related electronics. A single charging unit is used for charging the channel-specific pulse capacitors. Each coil is connected to its own channel; the electronics can drive up to six coils simultaneously, although here we use only five of them because they suffice for adjusting the stimulated location along two dimensions and rotating the electric field maximum. The auxiliaries contain miscellaneous electronics required for the operation and safety of the device. The electronics are located inside a grounded metal enclosure.

**Figure 1.**
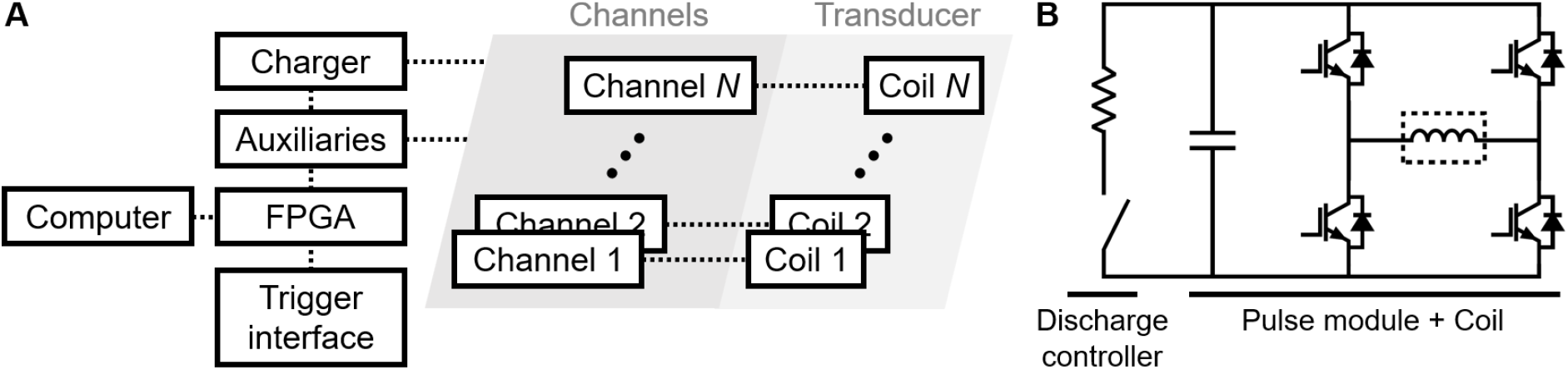
mTMS electronics. (**A**) An overview of the system modules. (**B**) A channel and a coil. The discharge controller is used to reduce the capacitor voltage, and the pulse module generates the stimulating current pulse through the coil (dashed box).

#### The control unit

is a field-programmable gate array (FPGA; PXIe-7820R; National Instruments, USA), interfaced through a custom-made LabVIEW (National Instruments) program in conjunction with specific logic-level trigger signals from external devices. The LabVIEW program, running on a dedicated computer, has an application programming interface, allowing one to develop components for the mTMS software in other programming environments.

#### The channels

form the core of the power electronics of the system; a channel comprises a pulse module and a discharge controller (Fig. 1B). A pulse module is composed of a high-voltage capacitor (E50.R34-105NT0, 1020 μF; Electronicon Kondensatoren GmbH, Germany) in parallel with a full-bridge circuit. Insulated-gate bipolar transistors (IGBTs; 5SNA 1500E330305; ABB Power Grids Switzerland Ltd., Switzerland) function as the switching elements in the bridge; due to the stray and load inductances present, they are protected by resistor–capacitor snubbing circuits (effectively 1 Ω and 1 μF in series). Custom-made driver boards control the switching of the IGBTs. Together with a coil, a pulse module forms the pulse-forming network of the stimulator. The discharge controller is a printed circuit board mounted directly on the pulse-capacitor terminals. The controller has two functions: first, it controls a discharge resistor (TE1000B1K0J, 1 kΩ, 1 kW; TE Connectivity, USA) parallel to the capacitor; second, a subcircuit on the board monitors the capacitor voltage and reports it to the control unit. Special attention was given to the physical layout of the power electronics to minimize the stray inductance in the pulse module.

#### The individual coils

of a multi-coil transducer are driven by separate channels. Thanks to the true parallelism of the FPGA, the current waveforms through all coils can be precisely controlled simultaneously. To avoid excessive circulating currents in the bridge after a stimulation pulse, the coils are characterized before the transducer is applied for brain stimulation, and the coil-specific waveforms are tuned so that no current is left circulating in the system after a pulse.

#### The charging

unit consists of a high-voltage charger (CCPF-1500; Lumina Power, Inc., USA) connected to a solid-state switching array, providing separate connections to all capacitors, one at a time. The maximum voltage is 1500 V, and the maximum charging time is about 700 ms per capacitor. The monophasic pulse waveforms used in this study (with a 60-μs rise time, a 30-μs hold period, and a 36.6–43.3-μs fall time [20]) reduce the capacitor voltage approximately 5–7% depending on the coil parameters; thus, a capacitor can be recharged to the voltage it had before a pulse in less than 100 ms.

#### The auxiliaries

contain various electronics modules that are vital for the proper operation of the mTMS device. Digital temperature sensors (DS18B20; Maxim Integrated, Inc., USA) serve the dual purpose of providing unique sensor identifiers to detect to which connector a particular coil is connected to, while also reporting the transducer temperature. Additionally, a sudden absence of a sensor reply can be utilized to detect a missing coil or a loose connection and to initiate an emergency shutdown. Most of the communication to the electronics on the high-voltage side is done via a communications interface that converts the electrical signals to and from the control unit into optical ones. Optical signaling provides a layer of isolation while also having excellent noise characteristics. Other circuit boards in this category include a power distribution module that delivers the required direct current power to the stimulator electronics, and an optically isolated trigger board that provides an interface for external triggering.

### Device operation

The operation of the mTMS device is based on forced current feed through the transducer coils, which is achieved by manipulating the electrical topologies of the coil-specific bridge circuits; see Fig. 2 [16,18,19,21,22]. Depending on the states of the IGBTs, a bridge circuit either connects its respective pulse capacitor in series with the coil connected to the channel (Fig. 2A,C), resulting in a damped oscillator circuit, or cuts the capacitor completely out of the circuit while also short-circuiting the coil’s ends (Fig. 2B). Even though the capacitor–coil circuit is oscillatory in nature, the duration we keep the capacitor connected to the coil is very short (tens of microseconds) compared to the oscillation period of the circuit (in the millisecond scale). The resulting current ramps are thus nearly linear.

**Figure 2.**
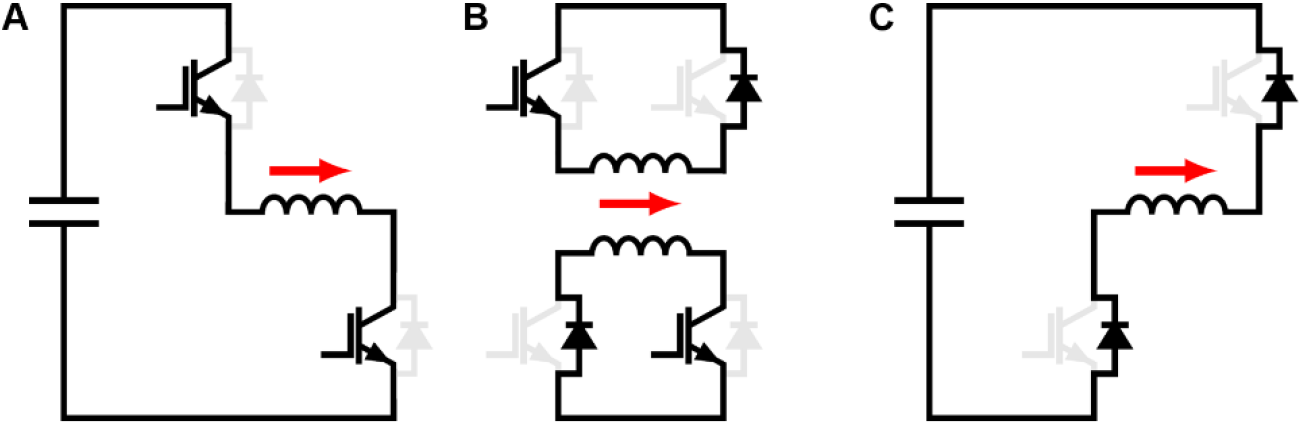
H-bridge operation. Depending on the states of the IGBTs, the H-bridge assumes one of the illustrated topologies. The red arrow indicates the direction of the coil current. For the sake of brevity, only states with positive coil current are shown. (**A, C**) The pulse capacitor is connected in series with the coil, allowing a change in the current through the coil. Current flows either through two IGBTs (A) or two flyback diodes (C). One IGBT from each side of the coil is conducting. (**B**) Coil-short configurations. The current through the coil keeps circulating through one IGBT and one flyback diode. Both IGBTs connected to one pole of the capacitor are conducting.

The capacitor–coil series configuration (Fig. 2A,C) leads to a changing current in the coil, a correspondingly changing magnetic field and an induced E-field in the brain. The coil-short configuration (Fig. 2B), on the other hand, leads to the current already flowing through the coil continuing its circulation, experiencing a slight decay mostly due to the resistance of the coil. The induced E-field due to this relatively slow change of current and magnetic field is negligible.

By incorporating multiple coils in a single transducer, the superposition of the E-fields is exploited to manipulate the spatial pattern, intensity, and direction of the E-field induced in the cortex [9]. The polarity and intensity of the E-field pattern from each coil can be manipulated by adjusting the rate of change of the current; this rate is proportional to the capacitor voltage in the corresponding channel. The total E-field induced in the cortex is the vector sum of the E-fields produced by the individual coils [9,13].

### mTMS transducer

We designed and built a 5-coil transducer to control the location and orientation of the peak of the induced E-field in a 30-mm-diameter cortical region (see Fig. 3 for an illustration of the transducer design geometry). The transducer design follows the 5-coil concept we introduced in [9]. The coil winding paths were generated with a minimum-energy optimization method [9] that utilized the interior-point method [23] and that was implemented in MATLAB 2020a (The MathWorks, Inc., USA). First, we modeled a commercial figure-of-eight coil (17 cm × 10 cm; Nexstim Plc, Finland) as 2,568 magnetic dipoles on a planar surface placed 15 mm away from the cortical surface (here modeled as a 7-cm-radius sphere using a 2,562-vertex triangular mesh; see Fig. 3) [24,25]. With the model of the commercial coil, we computed the induced E-field distribution for 8,964 coil placements (747 coil positions with 1-mm steps, 12 orientations with 30° steps in each position) with the maxima of the E-fields covering a 30-mm-diameter cortical region (Fig. 3). For each of these E-fields, we computed the corresponding minimum-energy surface current density in an octagonal plane section (30-cm diameter; 961-vertex triangular mesh) that would induce an E-field distribution with similar focality and intensity [16]. The optimization was performed for five distances between the sections and the cortical surface (15 to 27 mm in steps of 3 mm; see Fig. 3) to account for the winding thickness and mechanical factors that affect the construction of the physical coil. For each distance, we decomposed the optimized surface current densities with singular value decomposition and extracted the first five components, explaining 87.6–97.7% of the total variance depending on the distance. For each distance, we picked one of the five components so that the coils with the fastest attenuation of the E-field (or highest spatial frequencies [26]) were closest to the head [9]. Finally, we obtained the coil winding paths by discretizing the surface current density of each component in isolines and connecting them in series [27]. The process resulted in two four-leaf-clover coils (10 turns in each wing) at the bottom, two figure-of-eight coils (12 turns in each wing) in the middle, and an oval coil (26 turns divided into two layers connected in series) at the top.

**Figure 3.**
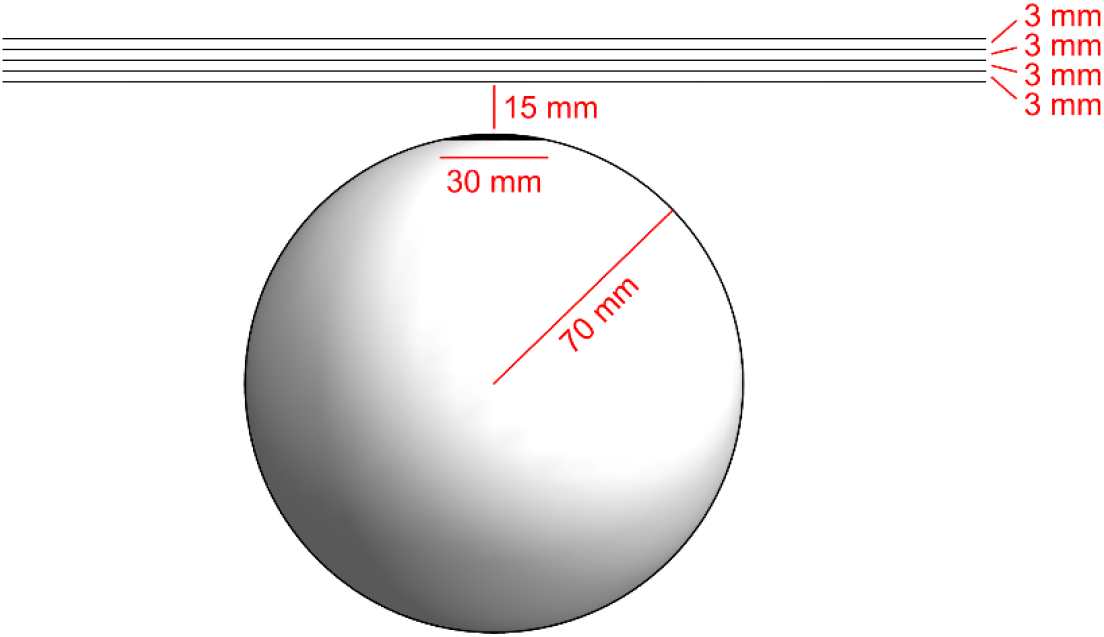
Transducer design geometry. In the optimization, the cortex was modeled as a 70-mm-diameter sphere. The black spherical zone on top of the sphere illustrates the 30-mm-diameter region within which the transducer was designed to be able to control the position and orientation of the E-field maximum. The coils were optimized on five 300-mm-diameter octagonal plane sections placed 15–27 mm above the cortex.

To manufacture the transducer, we designed five coil formers to accommodate the windings, a 5-mm-thick top cover to protect and insulate the wire connections, and a socket with a wooden rod attached to the top plate to ease transducer handling. The parts were designed in Fusion 360 (Autodesk, Inc., USA). The coil-former thicknesses were 4.0 mm (including a 1.0-mm-thick bottom) for the bottom-most coil, 3.9 mm (0.5 mm) for the top-most coil, and 3.5 mm (0.5 mm) for the coils in-between. The bottom thickness corresponds to the material thickness below the wire grooves. All parts were printed by selective laser sintering of 40% glass-filled polyamide (Proto Labs, Ltd., UK). Each coil was wound with copper litz wire (1.7-mm diameter; 3-layer Mylar coating; Rudolf Pack GmbH & Co. KG, Germany) in the grooves of the coil former and crimped to the end of a low-inductance TMS coil cable (Nexstim Plc). The assembly was potted and glued with epoxy for further mechanical strength and safety.

To characterize the manufactured 5-coil mTMS transducer, we measured the spatial distribution of the induced E-field, the self-inductance, and the resistance of each coil. The E-field distribution was sampled at 1,000 locations with our TMS characterizer [28], which gives E-field values on a 70-mm-radius cortex in a spherical head model. The center of the transducer bottom was at 85 mm from the center of the spherical head model. The self-inductance was measured with an LCR meter (1-kHz reference frequency; ELC-130; Escort Instruments Corp., Taiwan) and the resistance with a 4-wire measurement set-up using a bench multimeter (HP 34401A; Hewlett-Packard Co., USA). Given the realized coil winding paths, we also calculated the mutual inductances between the coil pairs with formulas of [29] implemented in Mathematica 12.3 (Wolfram Research Inc., USA). The duration of the pulse waveform was customized for each coil based on measurements with a Rogowski probe (CWT 60B; Power Electronic Measurements Ltd, UK) connected to an oscilloscope (InfiniiVision MSOX3034T; Keysight, USA) to ensure that no current was left circulating in the system after a pulse.

### Algorithm for electronic targeting

We applied the following algorithm to target the E-field in the cortex with the 5-coil transducer. In computing the E-field, the cortex and head conductivity boundaries were represented by triangular meshes extracted from individual magnetic resonance images and the coils were modeled according to the realized winding paths (see Data analysis). First, we specified a target location 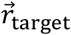 on the cortical surface and the desired E-field 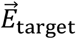 target at that location. We required that on the cortex (i.e., 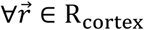, where 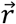 is the position and R_cortex_ the set of mesh nodes constituting the cortex), the E-field magnitude does not exceed 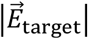. We searched for the coil currents **I** = [*I*_1_, *I*_2_, … , *I*_*N*_]^T^, where *I*_*i*_ is the current in the *i*th coil and *N* = 5 is the number of coils in the transducer, that minimize the magnetic energy *U* needed to induce the desired E-field pattern on the cortex. Given the inductance matrix

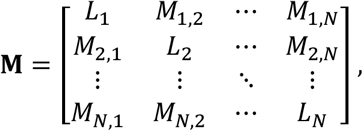

where *L*_*i*_ is the inductance of the *i*th coil and *M*_*i*,*j*_ the mutual inductance between coils *i* and *j* (in our case *M*_*i*,*j*_ ≈ 0 μH due to the designed approximate orthogonality of the coils: for ideal coils, the total magnetic flux through a coil due to any of the other coils would be zero), we can write the following formulation of the problem:

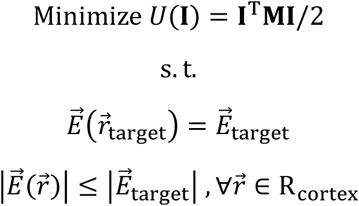

This optimization problem is similar to the ones we have encountered when designing optimal TMS coils [16,26]; thus, we solved it with the interior-point method [23]. We approximated each of the 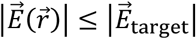 nodewise constraints with a convex constraint set. To keep the number of constraints small compared to the requirements of a 3-dimensional (3D) approximation [16], we applied an iterative approach in 2D. At each node, a 2D projection of the E-field was constrained to lie within a regular 16-gon, which provided a set of 16 linear constraints to restrict the norm of the projected E-field [26]. At the first iteration, we projected out the E-field along the direction of the node normal. In the subsequent iterations, we always started from the full 3D E-field and selected the projected-out direction to be perpendicular to the 3D E-field from the previous iteration. To reduce the number of constraints further, we constrained the E-field only at a downsampled set of those nodes in which its amplitude after the previous iteration exceeded 0.975 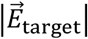 and a fixed set of four nodes around the target area. At each iteration, we appended the node positions at which 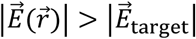 to the set R_max_, which was initialized before the first iteration as an empty set. We considered the optimization converged when the size of R_max_ did not increase.

To account for differences in the pulse waveforms due to coil-specific inductances and resistances, we scaled the obtained solutions (i.e., the applied capacitor voltages) so that the average E-field over the rising part of the monophasic pulses corresponded to the optimized **I**.

### mTMS motor mapping

To demonstrate mTMS in practice, we conducted a study on a 36-year-old healthy right-handed volunteer who provided written informed consent prior to his participation. The study was approved by an ethical committee of the Hospital District of Helsinki and Uusimaa and carried out in accordance with the Declaration of Helsinki.

Prior to the TMS experiments, we acquired structural magnetic resonance images (MRIs) of the subject’s head with a 3-T Magnetom Skyra scanner with a 32-channel receiver coil (Siemens Healthcare GmbH, Germany). For online neuronavigation, we acquired a T1-weighted image (cubic 1-mm^3^ voxels) with a magnetization-prepared rapid gradient-echo sequence. For E-field modeling with the boundary element method (BEM), we acquired a T1-weighted image with fat suppression and a T2-weighted image (both with cubic 1-mm^3^ voxels) [30].

In the TMS session, the participant sat in a chair and was instructed to keep his right hand relaxed. Surface electromyography (EMG) was recorded from the abductor pollicis brevis (APB), first dorsal interosseous (FDI), and abductor digiti minimi (ADM) muscles of the right hand with an EMG device (500-Hz low-pass filtering, 3-kHz sampling frequency; Nexstim eXimia; Nexstim Plc) with the electrodes in a belly–tendon montage. TMS was administered with the 5-coil mTMS transducer driven by our mTMS electronics. The pulse waveforms were monophasic with a 60-μs rise time, a 30-μs hold period, and an appropriate fall time (36.6–43.3 μs depending on the coil; Fig. 4) [20]. The transducer placement with respect to the subject’s head was monitored with a Nexstim eXimia neuronavigation system (Nexstim Plc).

**Figure 4.**
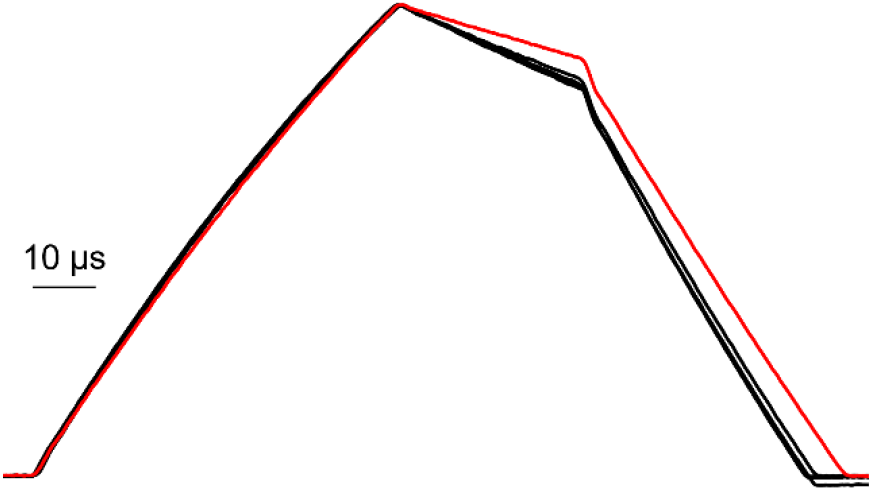
Measured current waveforms for the five coils at 1500 V. The red trace corresponds to the oval coil that has the lowest resistance and thus also the slowest decay of the coil current during the hold period; the other curves differ only slightly from each other. The data have been lowpass filtered at 1 MHz and normalized to the maximum of each curve.

First, with the bottom figure-of-eight coil and a fixed stimulator intensity, we searched manually for the direction and placement of the transducer leading to the largest motor evoked potentials (MEPs) in the APB (so-called APB hotspot). Then, we determined the resting motor threshold (RMT; 50% of the responses with a peak-to-peak amplitude exceeding 50 μV) of APB with a threshold tracking technique utilizing 20 stimuli at that target [31]. The ISI was randomized between 4 and 6 s.

To acquire a motor map, we kept the mTMS transducer fixed above the APB hotspot and adjusted the stimulation target electronically by varying the relative coil currents to mimic the movement of a figure-of-eight coil in a conventional mapping. We had predefined 100 target points on the left precentral gyrus and the desired E-field direction at each target; in the mapping, we aimed at 54 of these targets (i.e., those that were within the reach of the transducer) and the APB hotspot. The BEM-estimated induced E-field at the aimed location was kept at 110% RMT (i.e., 110% relative to the amplitude of the E-field maximum induced by the figure-of-eight coil at the RMT intensity) and its direction perpendicular to the precentral gyrus. The targets were stimulated in a pseudorandom order with an ISI of 4–6 s. We repeated the mapping 10 times.

### Data analysis

We computed the E-fields needed in the motor mapping experiment using a four-compartment volume conductor model and our surface integral solver [32]. We constructed the anatomical model from T1- and T2-weighted MRIs using the SimNIBS headreco pipeline [30]. We downsampled and smoothed the pial, skull, and scalp surface meshes, resulting in a boundary element mesh with a total of 33,592 vertices, of which 21,949 were on the pial boundary (3.5-mm mean vertex spacing on the pial surface). Using these surfaces and LGISA BEM solver [33], we built a four-compartment volume conductor model that contains the brain (conductivity 0.33 S/m), cerebrospinal fluid (1.79 S/m), skull (0.0066 S/m), and scalp (0.33 S/m).

As field computation space, we used a region of mid-cortical surface, which was represented with a dense mesh (11,811 vertices, 0.97-mm mean spacing) around the hand-knob area and with a 2-mm mesh elsewhere around the target region. The coil windings were exported from the design program as ordered point sets that formed polylines with 13,400–20,400 segments per coil describing the manufactured winding paths (series-connected isolines). These segments were further discretized using current dipoles, resulting in 13,600–20,800 dipoles per coil model. The E-field was computed in field space reciprocally using the volume conductor model and coil models otherwise as described in [32], but the coil integrals, i.e., the magnetic fluxes through the coils were computed using the circulation of the vector potential:

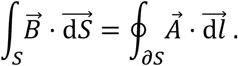

We split the EMG data into trials around the TMS pulses and subtracted the mean of the baseline signal at −100…0 ms from the whole trial. We rejected trials for which the absolute value of the EMG signal exceeded 20 μV within the 100 ms preceding the TMS pulse. For each accepted trial, we calculated the MEP amplitude as the peak-to-peak signal amplitude between 20 and 50 ms. Finally, for each stimulation target and muscle, we calculated the median MEP amplitude.

## Results

Figure 5A shows the 5-coil mTMS transducer placed on the scalp. Photos of the individual coils are shown in Fig. 5B. The measured E-field patterns are shown in Fig. 5C. The measured resistances and inductances of the coils were in the range of 79–109 mΩ and 15.2–17.1 μH, respectively. The calculated coupling coefficients describing the mutual inductances were 0–0.03 depending on the coil pair.

**Figure 5.**
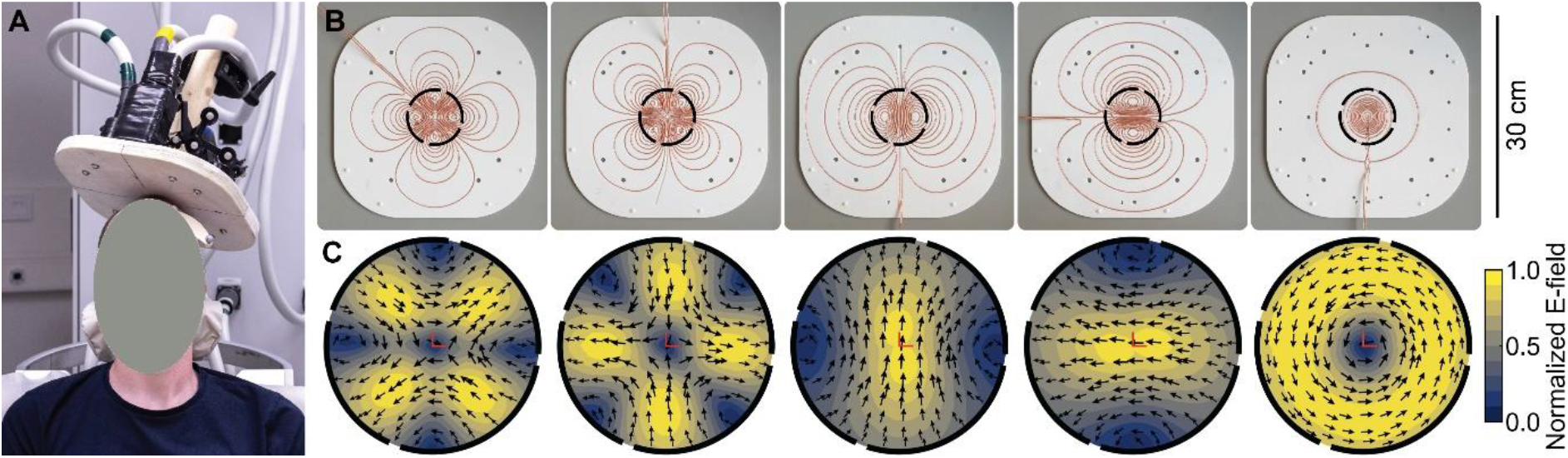
5-coil mTMS transducer. (**A**) The 5-coil mTMS transducer on the scalp. (**B**) Photos of the coil windings in the 3D-printed coil formers (30 cm × 30 cm). In the assembled transducer, the leftmost coil former is in contact with the scalp, and the rightmost coil former is the furthest from the scalp. (**C**) The E-field distribution of each coil in (B) measured with our robotic probe. The arrows indicate the E-field direction, the red lines mark the location below the transducer center, and the black circles (44-mm diameter) indicate the spatial extent of the E-field maps relative to the coils in (B).

The capacitor voltage at the RMT was 963 V, or 64% of the maximum (1500 V). At the RMT, the calculated time-averaged maximum E-field on the cortex during the rising part of the current waveform was 140 V/m at the APB hotspot. Figure 6 shows the calculated E-field at the APB hotspot and five examples of the optimized E-field targeting more lateral and medial locations of the precentral gyrus. The stimulation of the ABP hotspot was realized with the bottom figure-of-eight coil only. The other targets were reached by driving concurrently appropriate pulses through all five coils in the transducer. Figure 7 shows how the median MEP amplitude varied across the targeted primary motor cortex. We notice that for APB and FDI, we obtained large MEPs from an area that appeared more lateral to the region leading to the largest MEPs in the ADM. The maximum distance between the targeted points is 28 mm, which is on par with the 30 mm used as a design parameter for the available target region of the transducer.

**Figure 6.**
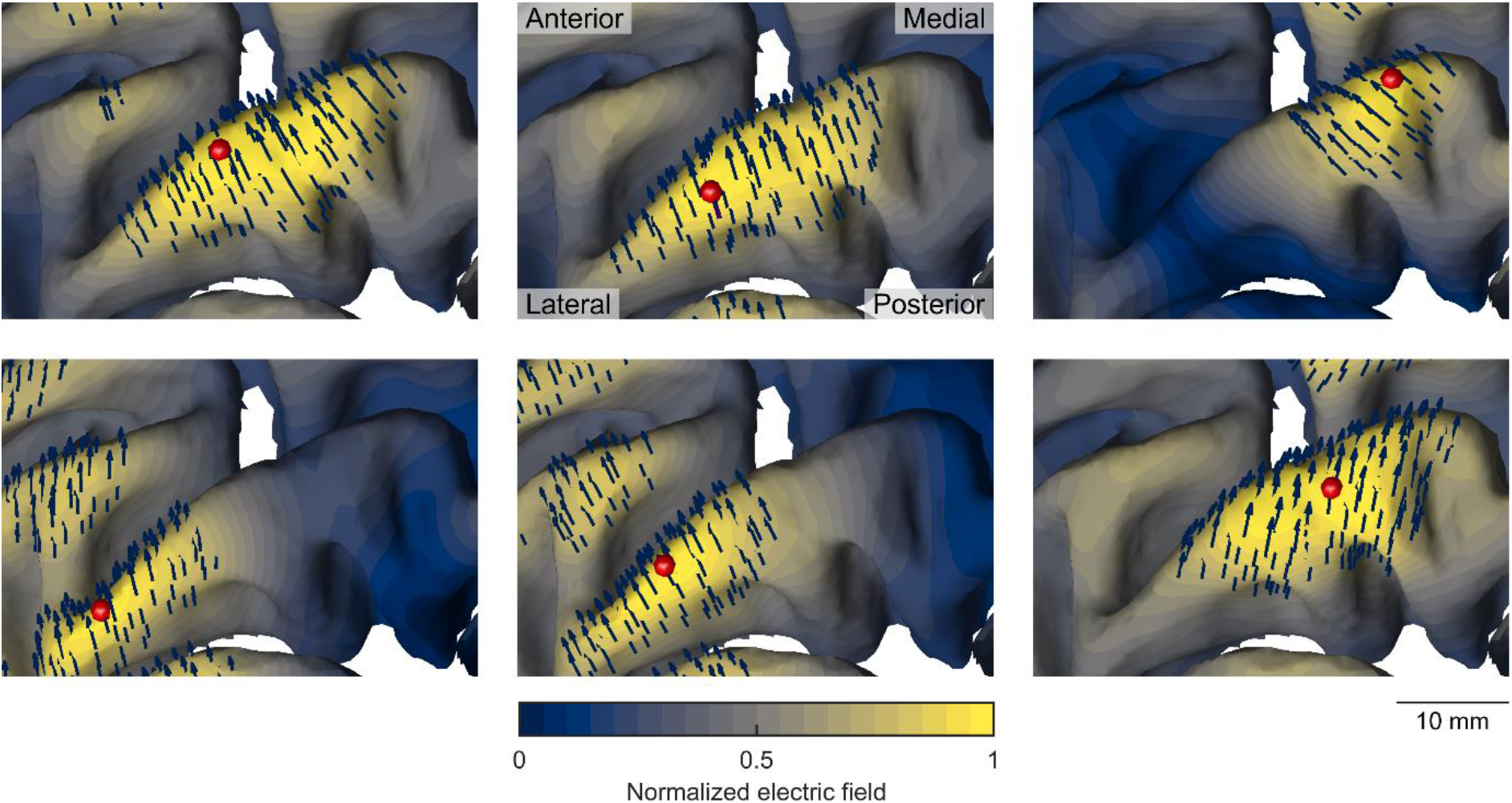
Examples of the induced E-fields. The estimated E-field at the APB hotspot (in the middle of the top row) and five examples of the optimized E-field with the same transducer placement but with the E-field maximum at translated targets. The red marker indicates the target location on the precentral gyrus, and the arrows show the E-field direction in the region where the E-field magnitude exceeds 70% of its maximum.

**Figure 7.**
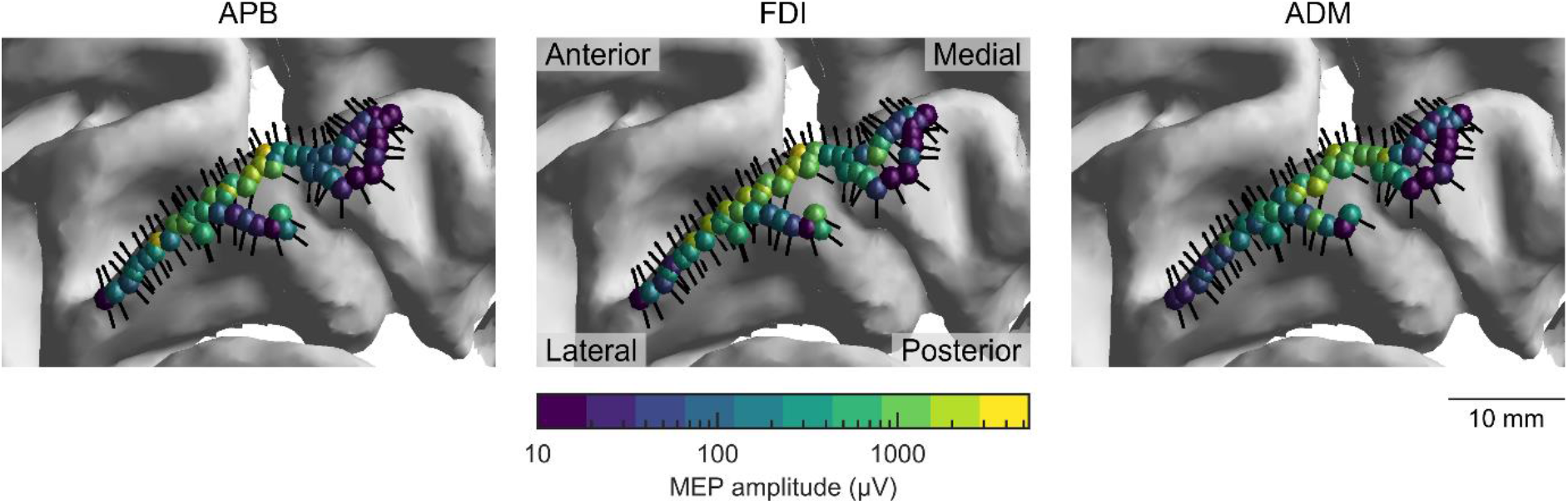
MEP responses. The marker color visualizes the median MEP amplitude for E-fields targeted to the marker location on the precentral gyrus in the orientation indicated by the associated black arrow. The results are shown separately for APB, FDI, and ADM.

## Discussion

Our mTMS system with its 5-coil transducer enables electronically targeted brain stimulation within a cortical region approximately 30 mm in diameter. The 5-coil transducer, which implements a design conceptually introduced in [9], allows automated mapping of the motor areas (Fig. 7), with the induced E-field oriented at will, as in our demonstration according to the gyral anatomy (Fig. 6) [34]. We were able to discern cortical motor representations, with results in agreement with earlier single-coil findings, showing that ADM is best activated with the E-field targeted more medially compared to FDI or APB [34,35].

The developed E-field targeting approach that employs convex optimization and BEM computations allows accurate adjustment of the induced E-field to target the desired cortical location. It also makes the entire mapping process easy and suitable for automation, as no manual transducer or coil movement is needed. Due to the convoluted cortical geometry, it may, however, be difficult to obtain the maximum E-field at some targets, e.g., those that are deeper than their surroundings [36]. This limitation applies, however, also to conventional TMS [36]. To minimize such problems, we manually selected points on the gyrus. For larger studies, it might be beneficial to develop a robust automated method to select the targets. The developed optimization formalism is quite general and can be expanded, e.g., to include constraints also for the coil currents to limit their amplitude (or the maximum rate of change) based on hardware limitations. Similarly, although in this study the only constraints on the E-field were the location and orientation of its maximum, other constraints may be added. For example, one may want to limit the E-field amplitude in specified non-targeted regions below a threshold or to have constraints on the E-field component perpendicular to the sulcal walls.

Figure 4 shows that there are minor differences between the current waveforms of the individual coils. In particular, the current decay in the oval coil is slightly slower than in the other coils. If we, however, assume that the neuronal activation occurs during the initial rising part of the current waveform [37– 39] or at the latest shortly after it [20], the differences in the decaying parts of the current waveforms have a negligible effect on the location of neuronal activation. The other minor differences of the waveforms were treated by considering the average E-field during the rising part as the effective stimulus strength, corresponding to the approximation that the neuronal strength–duration time constant (about 200 μs) [38] is much longer than the initial rising part of the pulse (60 μs). In this study, the weak mutual couplings between the coils (coupling coefficients on the order of 0–0.03) were neglected. One could, however, adjust the driving capacitor voltages to compensate for these couplings [9].

In the present mTMS system, we have implemented electronics capable of controlling up to six coils simultaneously. Since the mTMS electronics and the transducer are separate parts of the system, one can design special-purpose transducers without changing the electronics. For example, the 30-mm targeting range in the cortex can be made larger or E-field focality can be made adjustable. To expand the cortical region within which the E-field can be targeted beyond the 30-mm-diameter region demonstrated in this study, one may (1) develop a transducer with more than five coils [9,40], (2) reduce the desired E-field focality to design a five-coil transducer with a wider control region [40], or (3) implement a transducer that follows the head curvature [40]. To study interhemispheric communication in motor networks for the study of motor control [41], one may use the 5-coil transducer on one hemisphere while stimulating the contralateral hemisphere with a separate figure-of-eight coil. The 6th channel in our electronics will allow experimenting with such new designs in a flexible way.

In addition to automated cortical mapping, which may simplify, e.g., presurgical planning [42,43], mTMS will allow electronic stabilization to compensate for head movement during a TMS session faster than the existing robotic control [6]. mTMS also allows stimulating nearby targets with millisecond-scale interstimulus intervals [10], which may prove beneficial for developing new treatment and rehabilitation protocols. With physiological feedback from electroencephalography or electromyography recordings, mTMS enables closed-loop stimulation paradigms where stimulation targets are derived from the data gathered during the stimulation sequence [11,12].

## Conclusion

The developed mTMS system and the algorithm for E-field targeting enable electronically targeted TMS within a cortical region.

## Acknowledgements

This project has received funding from the Academy of Finland (Decisions No. 294625, 306845, and 327326), the Finnish Cultural Foundation, the Instrumentarium Science Foundation, the Jane and Aatos Erkko Foundation, and the European Research Council (ERC) under the European Union’s Horizon 2020 research and innovation programme (grant agreement No. 810377). The authors thank Matti Aho, Gustaf Järnefelt, Matti Mielonen, and Marko Ollikainen for useful discussions, Michael Serué for his work on the IGBT drivers, and Mikko Raskinen (Aalto University) and Tuomas Mutanen for the photos. The authors acknowledge the computational resources provided by the Aalto Science-IT project. The coil cables were donated by Nexstim Plc. Omnia helped with the bending of busbars for the pulse modules.

